# Proteome profiling of brain vessels in a mouse model of cerebrovascular pathology

**DOI:** 10.1101/2023.09.27.559361

**Authors:** Arsalan S. Haqqani, Zainab Mianoor, Alexandra T. Star, Flavie Detcheverry, Danica B. Stanimirovic, Edith Hamel, AmanPreet Badhwar

## Abstract

A cerebrovascular pathology that involves altered protein levels or signaling of the transforming growth factor beta (TGFβ) family has been associated with various forms of dementia, including Alzheimer disease (AD) and vascular cognitive impairment and dementia (VCID). Transgenic mice overexpressing TGFβ1 in the brain (TGF mice) recap VCID-associated cerebrovascular pathology and develop cognitive deficits in old age or when submitted to comorbid cardiovascular risk-factors for dementia. Here, we characterized the cerebrovascular proteome of TGF mice using mass-spectrometry (MS) based quantitative proteomics. Cerebral arteries were surgically removed from 6-month-old-TGF and wild-type mice, proteins extracted and analyzed by gel-free nanoLC-MS/MS. We identified 3,602 proteins in brain vessels, with 20 demonstrating robust altered levels in TGF mice. For total and/or differentially-expressed proteins (*p*≤0.01, ≥2-fold change), using multiple databases, we performed protein characterization, and identified proteins demonstrating RNA-transcripts in both mouse and human cerebrovascular cells, and known to be present in human-extracellular-vesicles (EVs). Dysregulated proteins point to perturbed brain vessel vasomotricity, remodeling, and inflammation. Given that blood-isolated EVs are novel, attractive and a minimally invasive biomarker discovery platform for the age-related dementias, several proteins identified in this study can potentially serve as VCID markers in humans.

## INTRODUCTION

With advancing age, blood vessels in the brain become increasingly vulnerable to pathologies. Damage to the brain’s vasculature disrupts the neurovascular unit, a multicellular system in which neurons, vascular cells (smooth muscle, pericytes and endothelial cells), astrocytes and microglia collaborate to ensure proper brain function [1,2]. Cerebrovascular damage can cause and/or aggravate age-related cognitive decline and dementia [3–5]. Affecting an estimated 55 million people worldwide, age-related dementias are a major cause of disability and dependency among the elderly (<u>World Health</u> <u>Organization</u>). A cerebrovascular pathology, partly involving altered signaling or increased levels of the cytokine transforming growth factor beta 1 (TGFβ1), has been associated with various types of dementia, including the two most frequent forms of dementia, namely Alzheimer’s disease (AD) and vascular cognitive impairment and dementia (VCID) [6–10]. Additionally, both hereditary small vessel diseases with cognitive deficits, cerebral autosomal dominant arteriopathy with subcortical infarcts and leukoencephalopathy (CADASIL) and cerebral autosomal recessive arteriopathy with subcortical infarcts and leukoencephalopathy (CARASIL), show loss of function of the high temperature requirement A serine peptidase 1 (HTRA1) gene, which results in upregulation of TGFβ signaling [11,12]. In particular, immunohistochemical analysis of the cerebral small arteries in CARASIL patients show increased TGFβ1 protein expression in the tunica media [11]. Alterations in TGFβ family signaling forms the basis for several vascular disorders in human, including hereditary hemorrhagic telangiectasia [13–16] and primary pulmonary hypertension [17]. Importantly, accumulating lines of evidence point to a link between cerebrovascular health and the TGFβ family signaling [6,7]. Increased risk of sporadic vascular dementia has been associated with the Pro10Leu single nucleotide polymorphism (SNP) in the *TGFβ1* gene [8], which is known to impact TGFβ1 protein level [9].

TGFβ family members, including TGFβ1, are multifunctional cytokines that bind to type I, II and III receptors [18]. Mouse models lacking TGFβ signaling components such as type I (e.g., TGFβRII) or II (e.g., ACVRL1, TGFβRI) receptors die mid-gestation due to impaired vascular development [19]. Transgenic mice overexpressing a constitutively active form of TGFβ1 in the brain (TGF mice), originally developed to recapitulate the cerebrovascular pathology seen in AD [20], were later found to lack the cerebral amyloid angiopathy typical of AD [21,22] and, as such, were found to better recap the cerebrovascular pathology seen in VCID. Cerebrovascular structural abnormalities observed in TGF mice include thickened vascular wall, microvascular injury and degeneration, such as smaller capillary endothelial cells and pericytes, and abnormal chromatin condensation in endothelial cell nuclei; and string vessel pathology [20,21,23,24]. Functional abnormalities such as impaired vascular reactivity primarily related to endothelial-mediated dilatation, chronic cerebral hypoperfusion, and compromised neurovascular coupling are also landmarks of TGF mice [23,24]. While TGF mice either do not develop cognitive deficits or only in advanced age [25,26], they readily do so when submitted to a comorbid cardiovascular risk factor for dementia, hence being a good model of VCID [27–29].

In recent years, -omics approaches, such as single-cell/nucleus transcriptomics and proteomics, have been used to comprehensively profile the mammalian cerebrovasculature [30–33]. These approaches have provided insight into transcriptome- and proteome-level changes in brain vessels from AD patients and mouse models [34,35]. However, to date, VCID-related pathology of brain vessels lacks similar profiling. Addressing this gap in knowledge, we set out to (a) characterize the cerebrovascular proteome of TGF mice using mass spectrometry (MS)-based quantitative proteomics, as well as (b) identify proteins with biomarker potential in human using an *in silico* bioinformatics approach. Given that proteins are regarded as effectors of biological functions, proteomics findings are generally considered directly suited for biomarker and drug development work.

## MATERIALS AND METHODS

### Mice

Six month-old transgenic TGF mice and their C57BL/6J wild-type (WT) littermates were used in this study. TGF mice overexpress a constitutively active form of TGFβ1 under the control of the glial fibrillary acidic protein (GFAP) promoter on a C57BL/6J background (line T64) [36]. To eliminate sex-related differences in brain structures [37] and potentially in the vasculature [38], only males were used. A total of 18 mice, in particular, 9 TGF mice and 9 WT littermates, were employed based on expected variances and differences between groups by our previous and other studies [12,23,35]. Mice were housed under a 12-hour light-dark cycle, in a room with controlled temperature (23°C) and humidity (50%). Mice had access to tap water and food (Teklad Rodent chow, Research Diets Inc., New Brunswick, NJ, USA) ad libitum. All experiments were performed in compliance with the Animal Ethics Committee of the Montreal Neurological Institute and the Canadian Council on Animal Care guidelines, and complied with the ARRIVE 2.0 guidelines [39].

### Surgical extraction of cerebral arteries

The circle of Willis and major cerebral arteries along with their main branches were surgically removed from mice and individually stripped from the attached pia matter to obtain a clean preparation of vascular tissue, as previously described [33]. Arteries extracted from three mice were combined to constitute one biological replicate and stored at -80°C. Three biological replicates (B1, B2, B3) were prepared for each of the two groups.

### Cerebrovascular proteomics workflow

Protein extraction from mouse cerebral arteries was performed using a published and validated protocol [33]. Thereafter, processing of samples for MS analysis, as well as the MS runs, preprocessing, and bioinformatics were performed without knowledge of group allocation (i.e., WT or TGF). A brief description of the procedure is provided in Supplemental information.

### Generation of statistically significant protein lists

Statistics were performed with knowledge of group allocations using GraphPad Prism version 9.4.1. Specifically, proteins with altered levels between WT and TGF cerebral arteries were identified through parametric Student *t* and non-parametric Mann-Whitney U tests. The *p*-values were corrected for multiple testing using Holm-Šídák method. Varying stringency was used to generate a database of two lists of proteins with altered levels between WT and TGF cerebral arteries. List 1 proteins were as follows: only proteins with *p*<0.01, peptide score ≥35 (<1% false discovery rate), and fold change ≥2 (up or down) were included, whereas peptides showing high variability (>55%) among replicates were excluded. List 2, more stringent, consisted of List 1 proteins identified with >2 peptides.

### Characterization of proteins

The gene associated with each identified protein was queried using the UniProt database [40] using the UniProtKB accession number of each protein. Cellular Component Analysis and PANTHER Overrepresentation Test were conducted using the PANTHER Classification System (PANTHER version 17.0 Released 2022-02-22, accessed April 24 and 26, 2022, respectively) [41,42] on the UniProtKB accession number of identified proteins, and with *Mus musculus* selected for organism. For the Cellular Component Analysis, the PANTHER GO-slim ontology annotation-set was used, which contains 3,361 terms – namely, 2,267 biological process, 550 molecular function and 544 cellular component terms (http://www.pantherdb.org/panther/goSlim.jsp). Fisher’s Exact test followed by False Discovery Rate (FDR) correction, was used for PANTHER pathways overrepresented in genes associated with identified proteins in mouse brain vessels compared to the entire *Mus musculus* genome. The PANTHER pathway database consists of 177, mainly regulatory, signaling pathways. Protein interactors of TGFβ1 were identified by querying BioGrid database, IntAct Molecular Interaction database (http://www.ebi.ac.uk/intact/) [43], and an in-house database [35] that is a compendium of several databases, including BIND, BioGRID, HPRD, HIMAP, and EcoCyc databases. Protein interactions were restricted to those observed in *Mus musculus* and/or *Homo sapiens*.

### Demonstrating RNA-transcript presence of identified proteins in brain vasculature *Using public databases*

We analyzed published single-cell transcriptomics datasets to demonstrate that RNA transcripts of proteins identified were also present in mouse and/or human brain vascular cells. Analyses were performed on List 2 proteins only. For relative gene expression levels of proteins in mouse brain vascular cell types (e.g., endothelial, smooth muscle cells), two mouse cerebrovascular single-cell transcriptomics datasets [31,44] were downloaded from the NCBI GEO repository [45], and each gene ranked by its relative abundance. For correlation between mouse and human relative gene expression in brain vascular endothelial cells, we used the dataset compiled by Yang and colleagues [34].

### Demonstrating presence of identified proteins in human extracellular vesicles (EVs) *Using public EV database*

Vesiclepedia [46], an extracellular vesicle (EV) database version 5.1, was used to identify List 1 and 2 proteins detected in the TGF cerebrovasculature and also detected in human EVs. These proteins constitute a rich source of brain vasculature-specific biomarkers, as well as receptors known for delivering molecules across the blood-brain-barrier.

### By conducting proteomics of plasma and plasma EVs

Five hundred (500) µL of purchased human plasma (from BioIVT, Westbury, NY; *bioivt.com*) was precleared by centrifugations at 1,500 ×g for 10 minutes and at 10,000 ×g for 10 min. The precleared plasma (supernatant) was used for abundant protein depletion or total EV isolation. For depletion of abundant proteins, 10 µL of precleared plasma was loaded onto High-Select™ Top14 Abundant Protein Depletion spin columns (ThermoFisher, Waltham, MA; catalog # A36369) and flowthrough collected as per manufacturer’s instructions. For isolation of total EVs, 150 μL of precleared plasma was loaded onto qEVsingle 35 nm size exclusion chromatography columns (IZON, Westbury, MA; product code # ICS-35) and total EVs were isolated as per manufacturer’s instructions. The precleared plasma, depleted plasma, and total EV fractions (200 µL of fractions 6-8) were analyzed by proteomics as recently described [47].

### Formatting guideline used for genes and proteins

For ease of reading, we have used the gene symbol for proteins. In accordance with the organism-specific formatting guidelines, mouse proteins (indicated by their gene symbols as mentioned above) have been the first letter in upper-case (e.g., Acta2), while human proteins have been fully capitalized (e.g., ACTA2). When indicating specifically a gene, the symbols have been italicized (e.g., *Acta2*, *ACTA2*).

## RESULTS

### Characterization of proteins detected in mouse brain vessels

We identified 3,602 proteins from high quality peptides (peptide score ≥35) in brain vessels of WT and TGF mice. Proteins identified included canonical vascular proteins, such Claudin-5 (Cldn5), Solute carrier family 2 (Slc2a1), and von Willebrand factor (Vwf). Using the UniProt database it was determined that the majority of proteins identified (N=3,575; 99.3%) were products of known genes. Cellular Component Analysis, conducted using the PANTHER Classification System, found 2,785 component hits in two main categories: cellular (N=2,167 genes) and/or protein-containing complex (N=618 genes). The top 5 cellular components were intracellular anatomical structures (N=1,624 genes), membrane (N=1,495 genes), organelle (N=1,290 genes), cytoplasm (N=1,037 genes), and cell periphery (N=754 genes), the latter defined as part of a cell encompassing the (a) cell cortex – cell region just beneath the plasma membrane and often containing a network of actin filaments and associated proteins, (b) plasma membrane, and (c) any external encapsulating structures (Fig. 1a). The top 5 protein-containing complex components were catalytic (N=181 genes), nuclear (N=157 genes), membrane (N=138 genes), ribonucleoprotein (N=96 genes), and intracellular (N=83 genes) (Fig. 1b).

**Figure 1:**
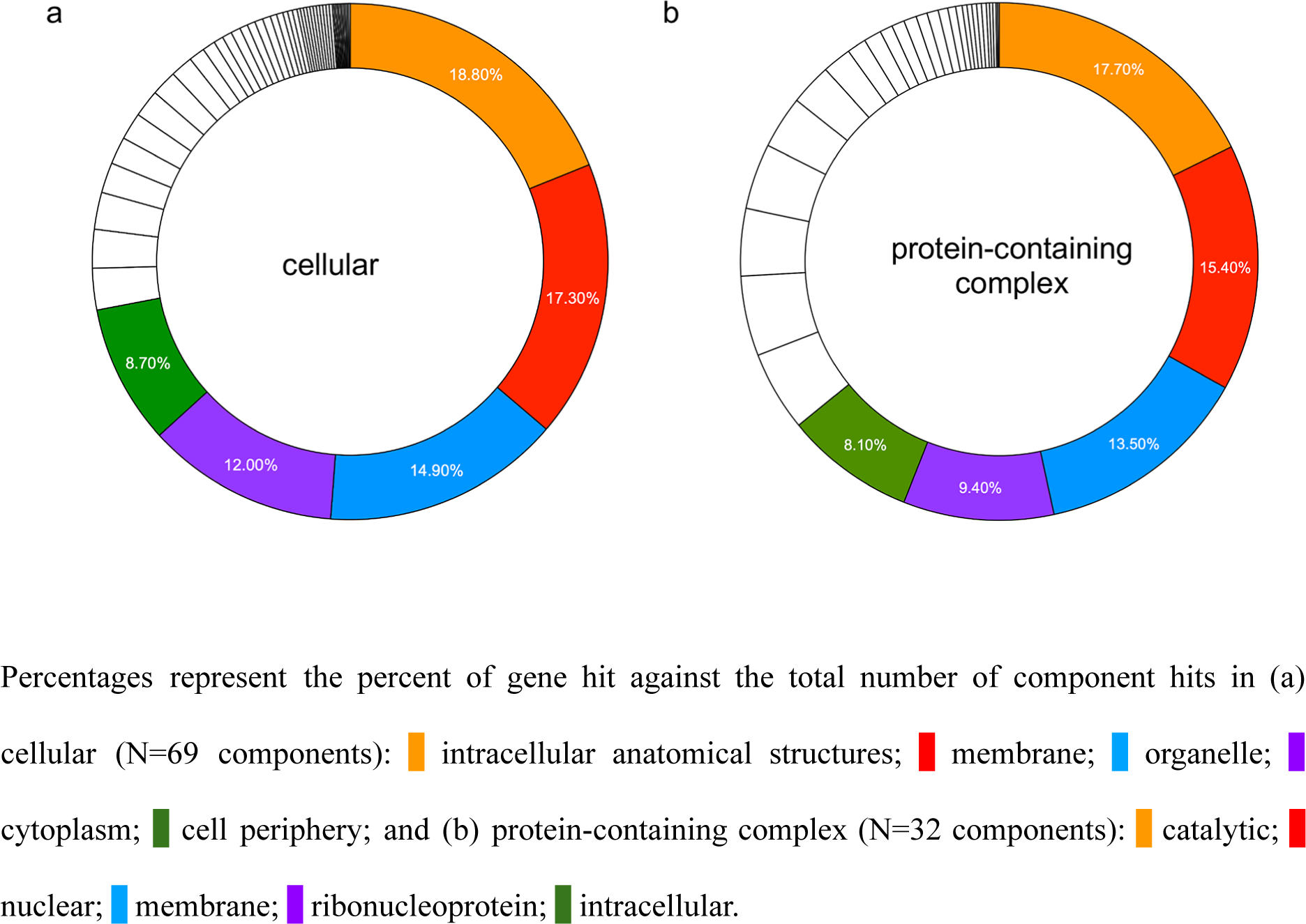
Percent distribution of the top 5 cellular and protein-containing complex components.

Using the PANTHER Overrepresentation Test, we identified seven pathways to be significantly (*p*<0.05, FDR corrected) overrepresented at a >2 fold enrichment (FEn) in mouse brain vessels, relative to the entire *Mus musculus* genome. These comprised pathways involved in (a) vasomotor regulation, namely, endothelin signaling (FEn=2.37) and 5-hydroxytryptamine/serotonin degradation (FEn=3.21) pathways, integrin signaling pathway (FEn=2.79), and cytoskeletal regulation by Rho GTPase (FEn=2.77), and (b) cellular metabolism/energetics, namely, tricarboxylic acid cycle (FEn=4.10), glycolysis (FEn=3.59) and insulin/IGF pathway-protein kinase B signaling cascade (FEn=2.53).

In our list of 3,602 proteins, we identified 102 (2.8%) direct interactors of TGFβ1, with ≥3 interactors detected from the following protein families: collagen (Col), heat shock proteins (Hsp), and large (Rpl) and small (Rps) ribosomal proteins (Fig. 2). We also identified 1,942 (53.9%) indirect interactors of TGFβ1.

**Figure 2:**
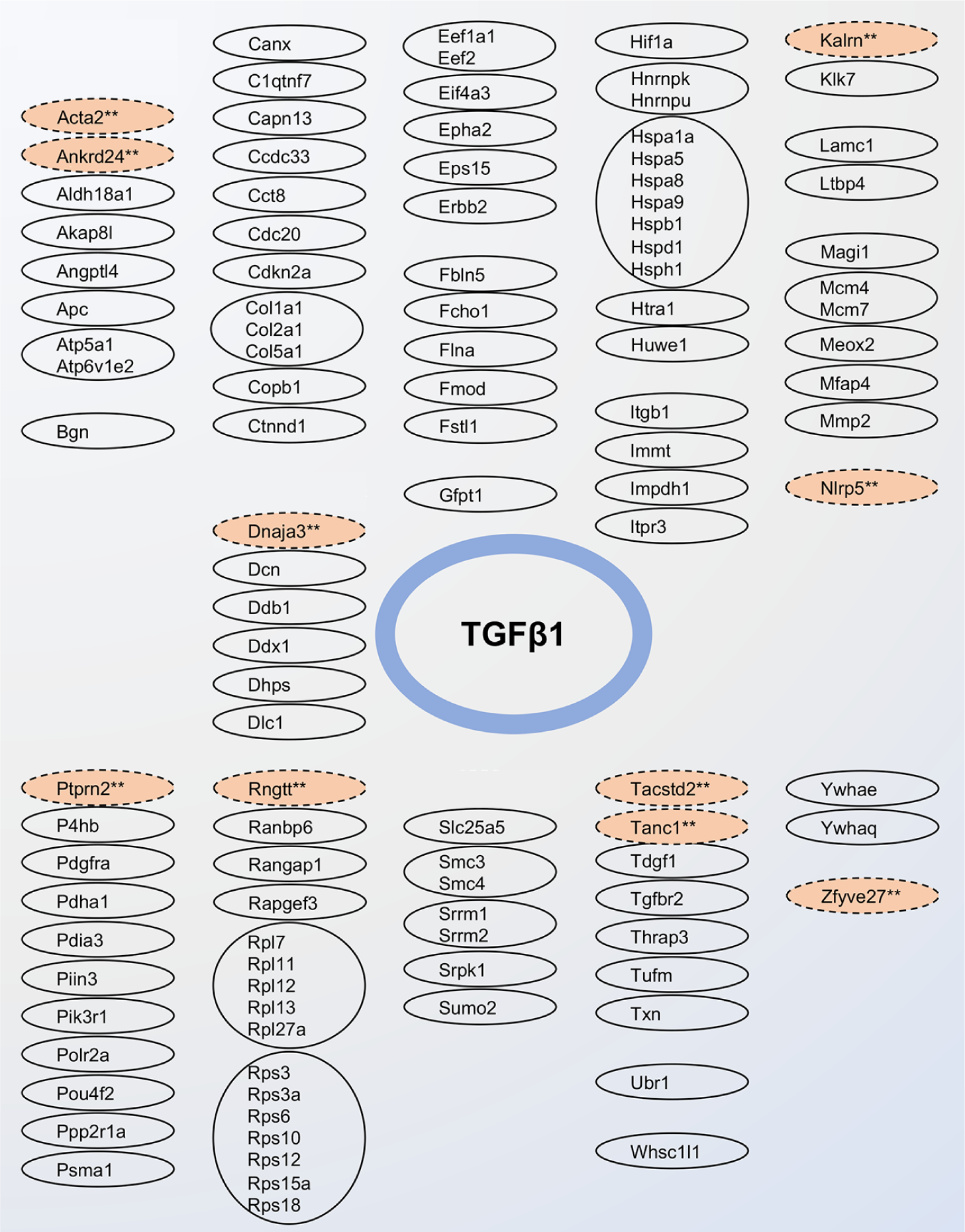
Protein interactors of TGFβ1. Solid circles indicate direct interactors, and dashed circles indicate indirect interactors. Protein interactors belonging to the same family of proteins are grouped together. Salmon colors depict TGFβ1 interactors with significant (*p*≤0.01) 2-fold level change in the cerebrovasculature of TGF mice.

### Proteins with altered levels in the brain vessels of TGF and WT mice

Compared to WT mice, 60 proteins (1.7%) showed significantly altered levels (*p*≤0.01 and 2-fold change) in the arteries of TGF mice. These 60 proteins constituted List 1 proteins (Supplemental Table 1). Of these, 20 proteins were significantly (*p*<0.01) detected with ≥2 peptides, with 10 showing level increases (Acta2, Ankrd24, Dll3, Adgrg2, Igdcc3, Kalrn, Nubpl, Ptprn2, Tanc1, Zfyve26), and 10 showing decreases (Ddx11, Dnaja3, Krt24, Nlrp5, Pof1b, Ptprd, Rngtt, Rp1l1, Tacstd2, Zfyve27) (Fig. 3, Supplemental Table 1). These 20 proteins constituted List 2 proteins (see full names of proteins in Table 1). Half (N=10) of List 2 proteins were indirect interactors of TGFβ1 (Fig. 2).

**Figure 3:**
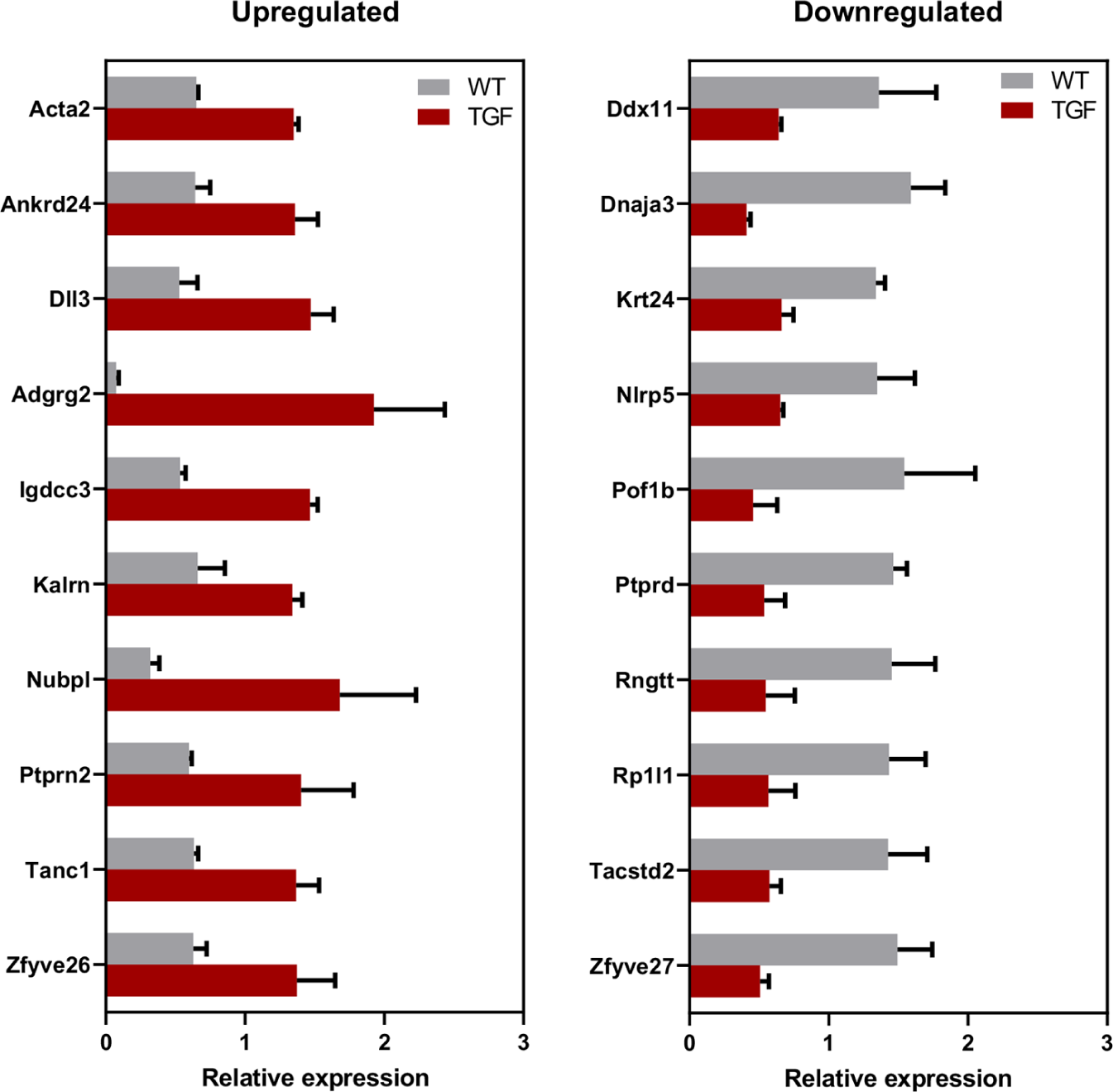
Relative expression of List 2 proteins in TGF brain vessels. Proteins with significantly altered levels in brain vessels of TGF mice relative to WT detected using the highest stringency in our database (*p*≤0.01, 2-fold change and ≥2 peptides) constitute List 2 proteins (N=20).

**Table 1:** Biomarker potential List 2 proteins detected in human extracellular vesicles. Note that the gene symbols of proteins are capitalized in accordance with the organism-specific formatting guidelines – in this case indicating human proteins. Abbreviations: p, protein; r, mRNA.

| ID<br>(human) | Common name | Vesiclepedia<br>Identified<br>molecule |  | Reported detected in<br>vascular cell and/or<br>blood, to date |  | Type |  |  |  |
| --- | --- | --- | --- | --- | --- | --- | --- | --- | --- |
|  |  | p | r | Vascular<br>[ref ID] | Blood<br>[ref ID] | Primary<br>HBEC | HBEC<br>EV | Plasma<br>(depleted) | Plasma<br>EV |
| ACTA2 | Actin Alpha 2 | X | X | Endothelial<br>[54,55] | Plasma<br>[51,52] | X | X | X | X |
| ANKRD24 | Ankyrin repeat domain-<br>containing protein 24 | X |  | - | - |  |  |  |  |
| DLL3 | Delta-Like protein 3 | X | X | - | - | X |  |  |  |
| ADGRG2<br>(GPR64) | Adhesion G Protein-Coupled Receptor G2 (G<br>Protein-Coupled Receptor<br>64) | X | X | - | - |  |  |  |  |
| IGDCC3 | Immunoglobulin<br>Superfamily DCC Subclass<br>Member 3 |  | X | - | - |  |  | X |  |
| KALRN | Kalirin RhoGEF Kinase | X | X | - | Plasma<br>[53] |  |  |  | X |
| NUBPL | Nucleotide Binding<br>Protein-Like | X | X | - | - |  |  |  |  |
| PTPRN2 | Receptor-type tyrosine-<br>protein phosphatase N2 |  | X | - | - |  |  |  |  |
| TANC1 | Tetratricopeptide Repeat,<br>Ankyrin Repeat, and<br>Coiled-Coil Containing 1 | X | X | - | - | X |  | X | X |
| ZFYVE26 | Zinc Finger FYVE-Type<br>Containing 26 | X | X | - | - |  |  |  | X |
| DDX11 | Probable ATP-dependent<br>RNA helicase DDX11 | X | X | - | - |  |  |  | X |
| DNAJA3 | DnaJ Heat Shock Protein<br>Family (Hsp40) Member<br>A3 | X | X | - | - |  |  |  |  |
| KRT24 | Keratin 24 | X | X | - | Plasma<br>[51] |  |  | X | X |
| NLRP5 | NOD-like receptor family<br>pyrin domain containing 5 |  |  | - | - | X |  |  |  |
| POF1B | POF1B actin binding<br>protein | X |  | - | - |  |  |  |  |
| PTPRD | Protein Tyrosine<br>Phosphatase Receptor<br>Type D | X |  | - | - | X |  |  | X |
| RNGTT | mRNA-capping enzyme | X | X | - | - |  |  |  |  |
| RP1L1 | Retinitis pigmentosa 1-like<br>1 |  |  | - | - | X |  |  |  |
| TACSTD2 | Tumor associated calcium<br>signal transducer 2 | X |  | Endothelial<br>[56] | - |  |  |  |  |
| ZFYVE27 | Protrudin | X | X | - | - |  |  |  |  |

### Comparison against published transcriptomics in mouse brain vascular cells

Examination of published single-cell transcriptomics data demonstrated that RNA transcripts of the above-mentioned 20 List 2 proteins are present in one or more cell types of the mouse brain vasculature. Abundance ranking of their gene expression in two separate single-cell transcriptomics databases, ranging from low (<30%) to very high (>90%) are shown in Fig. 4a and Supplemental Fig. S1. Comparison of transcripts in the arterial endothelial cell (aEC) and arterial smooth muscle cell (aSMC) categories identified 11/20 of these transcripts to be present in both cell types (Acta2, Ankrd24, Kalrn, Nubpl, Tanc1, Zfyve26, Ddx11, Dnaja3, Ptprd, Rngtt, Zfyve27) with an abundance ranking of ≥50% (indicating moderate to high abundance). In contrast, 1/20 transcript (Igdcc3), and 2/20 transcripts (Ptprn2, Tacstd2), demonstrated an abundance ranking of ≥50% in aEC and aSMC, respectively.

**Figure 4:**
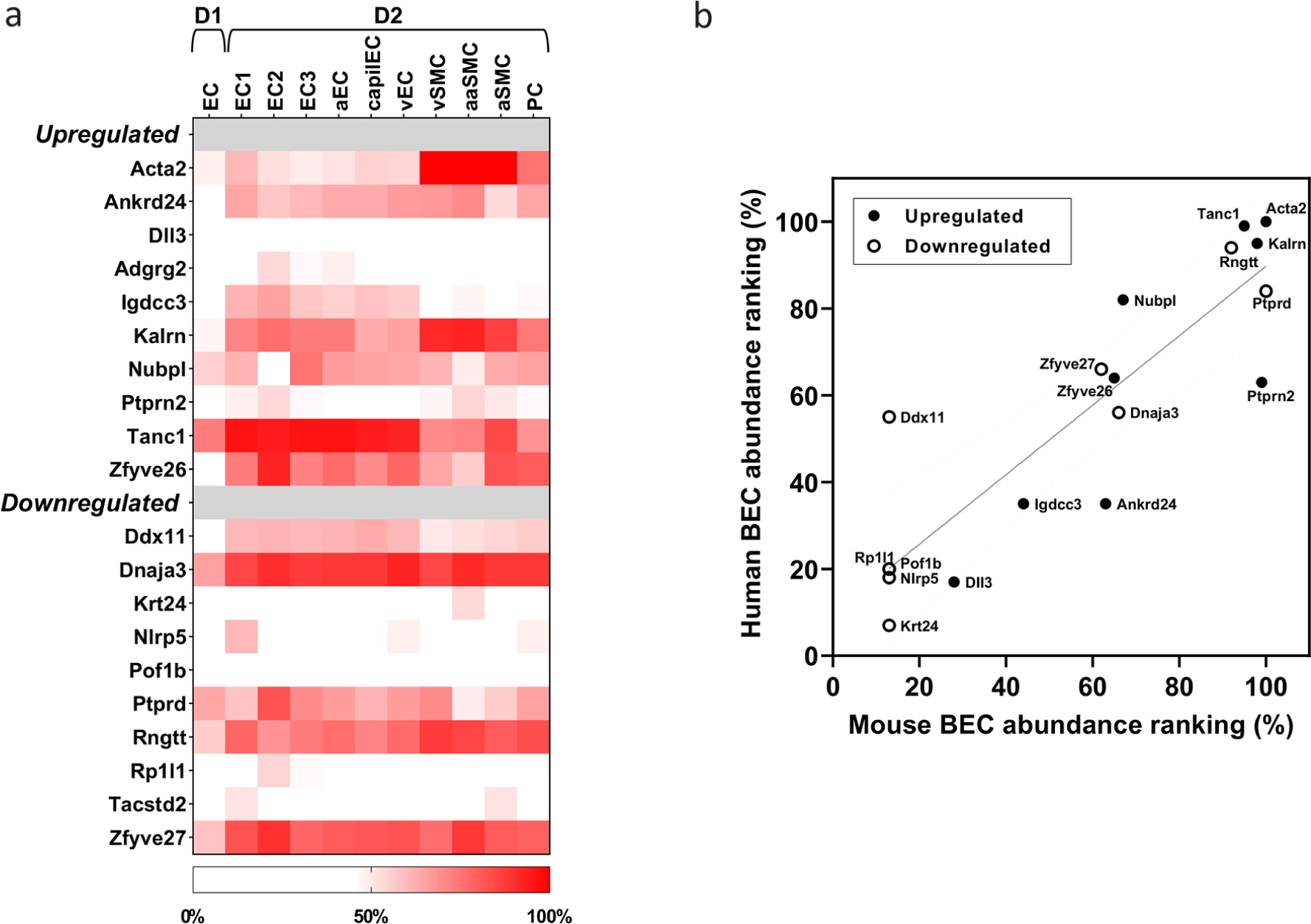
Relative gene expression of List 2 proteins in public vascular datasets. (a) Relative gene expression of the 20 List 2 proteins in mouse cerebrovascular single cell transcriptome databases D1 [44] and D2 [31]. Abbreviations: EC, endothelial cell; aEC, arterial EC; capilEC, capillary EC; vEC, venular EC; vSMC, venular smooth muscle cell; aaSMC, arteriole SMC; aSMC, arterial SMC; PC, pericyte; (b) Correlation between mouse and human relative gene expression of the 20 List 2 proteins in brain vascular endothelial cells. Upregulated and downregulated proteins are shown as closed and open symbols respectively. Solid line represents the linear regression (Pearson *r*^2^=0.64, *p*<0.001) and dotted lines are 99% confidence bands.

### Translatability of findings to human

To demonstrate the translatability of our findings (i.e., the differentially-expressed mouse proteins are also expressed in human brain vasculature), the relative gene expression of our List 2 proteins were compared in public RNAseq databases between human and mouse brain vascular endothelial cells. Most (N=17) of the proteins show similar gene expression in the two species as demonstrated by a strong correlation (Pearson *r*^2^=0.64, *p*<0.001) in the expression ranking between human and mouse (Fig. 4b). Additionally, the List 2 proteins were examined in proteomics datasets from human brain endothelial cells [48–50], which is part of the BBB Carta project [48]. Out of the 20 List 2 proteins, six were detectable in human brain endothelial cells, including ACTA2, DLL3, TANC1, NLRP5, PTPRD, and RP1L1.

### Blood biomarker potential in human

Using a publicly-available database, Vesiclepedia, we demonstrated that 80% (N=16) of the 20 List 2 differentially expressed proteins in the TGF cerebrovasculature have been detected in human EVs (Table 1), with several detected in easily accessible biofluids (e.g., plasma [51–53]). In addition, Acta2 and Tacstd2 were detected in EVs secreted by vascular endothelial cells [54–56]. Acta2 was also detected in EVs secreted by human blood-brain-barrier cells [49]. Overall, 70% (N=42) of the 60 TGF List 1 differentially expressed proteins (which includes List 2 proteins) were detected in human EVs (Supplemental Table 1).

In addition, the 20 proteins in List 2 were also validated in the proteome of healthy human plasma to demonstrate their biomarker potential. Depletion of abundant proteins from plasma was necessary to detect the presence of some of these proteins by proteomics. At least four of the List 2 proteins (ACTA2, KRT24, TANC1 and IGDCC3) were detectable in depleted human plasma (Table 1). An example of extracted ion-chromatogram and MS/MS spectrum for a TANC1 peptide in non-depleted and depleted plasma is shown in Supplemental Fig. S2 a and b.

Total EVs were also isolated from these plasma samples using size-exclusion chromatography and analyzed by proteomics. At least seven of the List 2 proteins (ACTA2, KALRN, TANC1, ZFYVE26, DDX11, KRT24 and PTPRD) were detectable in the total plasma EVs. Examples of extracted ion-chromatograms and MS/MS spectra for the peptides of ACTA2 and KALRN in plasma and total EVs are shown in Supplemental Fig. S3 a-d.

## DISCUSSION

TGF mice overexpress a constitutively active form of TGFβ1 in brain astrocytes and recapitulate the cerebrovascular pathology seen in VCID, and, except for the amyloid pathology, in AD. Particularly, they display thickened vascular walls due to accumulation of extracellular matrix proteins, string vessel pathology, impaired dilatory function, cerebral hypoperfusion, neurovascular uncoupling, and cerebral microhemorrhages [20,21,23,57,58]. Here, we demonstrate that the cerebrovascular proteome is significantly altered in TGF mice. Specifically, 60 proteins (1.7% of total identified proteins in WT mice) showed significant altered levels (*p*≤0.01 and 2-fold change) in the cerebral arteries of TGF mice compared to those of WT littermates. We focus our discussion on how level perturbations in List 2 proteins (N=20), identified using stringent criteria from the 60 List 1 proteins, relate to the cerebrovascular abnormalities seen in TGF mice, in particular perturbation in vasomotricity, remodeling, and inflammation.

### Perturbation in vascular tone and vasomotricity

We identified three proteins with altered levels in the TGF cerebral vasculature, namely, Acta2, Tacstd2, and Tanc1, that are known to influence vascular tone.

#### Increased Vascular Tone

The contraction of blood vessels in response to the stretch resulting from pulsatile blood flow is dependent on the interaction between thin and thick myofilaments, composed of Acta2 (or α-2 actin) and β-myosin, respectively [59,60]. Increased Acta2 level in the TGF cerebrovasculature likely increases vascular tone, hence contributing to the impaired vasomotricity observed in brain arteries of TGF mice [23]. Our finding of increased Acta2 levels in the TGF cerebral arteries parallels observations in peripheral arteries, where the overactivation of TGFβ1 signaling stimulates expression of smooth muscle contractile genes, including *ACTA2* [61,62]. Additional evidence linking Acta2 to the regulation of vascular tone include: (a) the observation that *Acta2*-null-mice demonstrate compromised vascular tone, contractility, and blood flow [63], and (b) the dilative phenotype found in *ACTA2* missense mutations seen in cerebral arteriopathy in human [64].

#### Attempt To Compensate By Promoting Vasodilation

In contrast to the vasocontractile phenotype promoted by upregulated Acta2, decreased Tacstd2 protein levels in the TGF cerebrovasculature point to an attempt at promoting vasodilation, via modulation of Ca^2+^ levels. Tacstd2 is known to increase Ca^2+^ levels in the cytosol, by inducing its release from internal stores [65,66], which increases vascular smooth muscle contractility [33]. Therefore, a decrease in Tacstd2 level would promote vasodilation. Tanc1, demonstrating increased levels in TGF brain vessels, is an indirect interactor of TGFβ1 [67], and a scaffold protein associated with regulation of excitatory (e.g., glutamatergic) synapse strength in both brain and non-brain tissue [68,69]. It is, therefore, possible that increased levels of Tanc1 is an attempt to promote blood vessel dilation via modulation of neurogenic signaling [70]. Indeed, Tanc1 has been shown to interact with NMDA (N-methyl-D-aspartate) and AMPA (α-amino-3-hydroxy-5-methyl-4-isoxazolepropionic acid) glutamate receptor subunits encoded by *GRIN2B* and *GRIA1* [71], and present in cerebral blood vessels [33,72].

In addition to the three proteins known to influence vascular tone, we also detected decreased levels of the ATP-dependent DNA helicase Ddx11, an alteration likely related to the contractile phenotype associated with chronic cerebral hypoperfusion in TGF mice [29,57,58]. Supporting this is the observation that in the vascular endothelium in human the *DDX11* gene is repressed under hypoxic conditions [73].

### Cerebrovascular remodeling

Remodeling of the cerebrovasculature is a key phenotypic feature of TGF mice [20,21,23,24]. In general, vascular remodeling is the process of altering blood vessel structure and arrangement via cell growth, proliferation, migration, death, and/or the production or degradation of the extracellular matrix [74]. Similar to the remodeling observed in TGF brain vessels, increase in TGFβ1 level is known to promote aspects of vascular modeling, such as a proliferative phenotype, in peripheral arteries, as demonstrated (a) in cultured vascular smooth muscle cells from human pulmonary artery [75], and (b) in murine carotid artery vascular media – due to localized overexpression of TGFβ1 in the endothelium [76]. We found that 11 of the 20 proteins with altered levels in the TFG cerebral vasculature were known influencers of vascular remodeling, namely: Dll3, Adgrg2, Krt24, Ptprd, Ptprn2, Igdcc3, Rngtt, Rp1l1, Tacstd2, Zfyve26, and Zfyve27. Below, we highlight their specific links with processes contributing to vascular remodeling.

#### Vascular Remodeling Via Promotion of Cell Proliferation

Recent literature suggests that TGFβ1 overexpression can upregulate the expression of several Notch signaling pathway proteins [77]. Indeed, an increase in Dll3 level, a core member of the canonical Notch signaling pathway, was observed in TGF vessels. Dll3 binds to the Notch family of receptors (Notch1 to 4) [78], of which Notch1 and Notch4 were detected in cerebral arteries of TGF and WT mice. In human, elevated DLL3 level has been linked with pathological angiogenesis via the DLL3/NOTCH4 signaling pathway [79]. Adgrg2, a G-protein coupled receptor, was increased in brain vessels of TGF mice. Literature indicates that G-protein coupled receptor signaling may play a role in arterial smooth muscle cell transformation [80]. In line with this, upregulation in *ADGRG2* transcripts has been reported during the transformation of quiescent human coronary artery smooth muscle cells to a proliferative and migratory phenotype observed in atherosclerosis [80]. Cytokeratin Krt24, a known anti-proliferative factor in the epidermis [81,82], was decreased in the TGF cerebrovasculature. Given that differential expression of cytokeratins has been observed in vascular smooth muscle cells, often in the setting of a proliferative state [83], decreased Ktr24 level likely contributes to cell proliferation and remodeling. Decreased Ptprd and increased Ptprn2 levels were also found in the TGF cerebrovasculature. These two proteins belong to the protein tyrosine phosphatase family of signaling molecules that regulate cellular processes, including differentiation and proliferation [84]. Decreased Ptprd protein level in peripheral artery (i.e., pulmonary, rodent) was found to alter vascular smooth muscle cell morphology and migration, primarily via focal adhesion and cell cytoskeleton modulation [85]. The protein Ptprn2 is a direct interactor of SMADs [86], the main downstream signal transducers for the TGFβ superfamily receptors. Variations in the *PTPRN2* gene have been associated with calcified atherosclerotic plaque, a subclinical marker of atherosclerosis, in human peripheral arteries [87,88], and familial stroke [89]. Increase in Ptprn2 (or human PTPRN2) protein level is also known to perturb the lipid-dependent sequestration of an actin-remodeling factor [90], a process capable of facilitating vascular remodeling. While the exact role of Igdcc3 (or human IGDCC3) protein in brain vessels is unknown, its gene expression pattern strongly overlaps with regions of high Wnt activity during embryonic development [91], suggesting that its increase in the TGF cerebrovasculature may be associated with a proliferative phenotype [92].

#### Attempt To Attenuate Vascular Remodeling

Levels of proteins Rngtt, Rp1l1, Tacstd2, Zfyve26, and Zfyve27, which are known to promote a proliferative phenotype in various cancers were decreased in the brain vasculature of TGF mice, likely as an attempt to attenuate vascular proliferation and remodeling. Reduction in Rngtt level, a protein that caps nascent mRNA [93], has been shown to compromise the activity of Wnt3a/β-catenin [94] and Hedgehog [95] pathways, both involved in a proliferative phenotype in the vessel wall [92,96]. Commonly associated with retinal degeneration, the *RPL11* gene has also been associated with brain arteriovenous malformations [97] and cancer [98,99] in human. Similarly, expression of the TGFβ1 response gene [100], *TACSTD2* (or *TROP2*), in cancer tissue (e.g., human glioma) has been shown to correlate with microvessel density, a commonly used marker for estimation of angiogenesis [101]. In addition, by conducting experiments in both human and mice, Guo et al. demonstrated that Tacstd2 (or human TACSTD2) promotes expression of MMP13 and PECAM1, two well-known angiogenesis factors, via activation of ERK-1 and -2 signaling pathway [102]. With regard to Zfyve26, decreased levels in mice is known to disrupt endolysosomal membrane trafficking [103], which in turn can exacerbate vascular calcification [104]. Therefore, its increase in the TGF cerebrovasculature may be a compensatory attempt at attenuating remodeling. Zfyve27 (or human ZFYVE27) was recently found to promote endothelial cell migration and angiogenesis in human, with its knockdown affecting these processes [105].

### Chronic Inflammation

While TGFβ1 has been known to function both as an anti- and a pro-inflammatory cytokine [106], a pro-inflammatory phenotype has been observed in TGF mice [24,107], including inflammation of brain vessels characterized by activated perivascular astrocytes and microglial cells [21]. TGFβ1 is known to stimulate the production of reactive oxygen species (ROS) in vascular cell types, including endothelial and smooth muscle cells [108,109]. ROS are free radicals derived from molecular oxygen, and their overproduction in the cerebrovasculature contributes to the inflammatory response [110] and promotes dysfunction by shifting cell bioenergetics [111]. Six of the 20 proteins with altered levels in the TFG cerebral vasculature, namely Nlrp5, Pof1b, Kalrn, Nubpl, Dnaja3, and Ankrd24, had links to inflammation, as described below.

#### Proteins Indicative of An Inflammatory Process

Decrease in Nlrp5 and Pof1b levels were observed in the TGF brain vessels. Nlrp5 (or human NLRP5) belongs to a class of cytoplasmic pattern-recognition receptors [112], and is known to be expressed in both mouse and human cerebrovasculature, largely in endothelial cells [31,113] and pericytes [31,114]. Given that treatment with hydrogen peroxide (an endogenous ROS) decreases the expression of NLRP5 in human cerebral endothelial cells, its decrease in the TGF cerebrovasculature may indicate ROS overproduction and an inflammatory environment [113]. With regard to Pof1b (or human POF1B), its gene expression in blood has been associated with ischemic stroke in human [115]. POF1B is also known to be highly expressed in epithelial cells with tight junctions, where it localizes at tight junctions and regulates cytoskeleton dynamics [116]. In the vasculature, *Pof1B*/*POF1B* expression has been observed in endothelial cells and pericytes [31,34,44], and may contribute to the regulation of the blood-brain-barrier permeability, similar to its function in the epithelium. Its decrease in the TGF cerebrovasculature may indicate a compromised blood-brain-barrier and an inflammatory environment.

#### Attempt At Reducing Inflammation

Furthermore, altered levels of the proteins Kalrn, Dnaja3, Nubpl, Spen and Ankrd24 may suggest compensatory attempts at reducing inflammation in the TGF cerebrovasculature. Increased Kalrn level was found in the TGF brain vessels. In human, KALRN, a guanine nucleotide exchange factor, is known to associate with and down-regulate inducible nitric oxide synthase (iNOS) activity, and thereby production of nitric oxide, a free radical [117,118]. Interestingly, polymorphisms in the *KALRN* gene have been associated with increased risk of both cerebrovascular and cardiovascular diseases, including ischemic stroke [119] and coronary artery disease [118,120,121]. Perturbed levels of the mitochondrial proteins Dnaja3 and Nubpl may also be an attempt at attenuating inflammation in the TGF cerebrovasculature via (a) decreased Dnaja3 associated dampening of innate immunity [122] and (b) increased Nubpl associated dampening of ROS production [123]. While little is known about Ankrd24, it can interact with nuclear factor kappa B (NF-κB) [43], a transcriptional activator of inflammatory mediators. Direct interaction of Ankrd24 with NF-κB may serve a similar function as the interaction of the stress response protein Ankrd2 with NF-κB repressor subunit p50, which results in potent repression of inflammatory responses [124]. Interestingly, vasomotor impairment in TGF cerebral arteries was restored by the nonsteroidal anti-inflammatory drug (NSAID) indomethacin and pioglitazone [107], the latter likely via activation of PPARγ receptors in brain vessels and reduction of NF-κB and IL-6 levels [125].

### Blood biomarker and translatability to human

In our study, we confirmed the presence of four List 2 proteins (ACTA2, KRT24, TANC1, and IGDCC3) in depleted human plasma and seven List 2 proteins (ACTA2, KALRN, TANC1, ZFYVE26, DDX11, KRT24, and PTPRD) in total plasma EVs. These findings hold significant promise for blood biomarker research, diagnostics, and treatment, particularly in the investigation of vascular brain injury in age-related dementias. Notably, a recent review [126] underscores the emerging role of EVs in blood plasma as biomarkers in age-related dementias, further emphasizing the importance of our results.

## Conclusion

By performing MS-based vascular proteome profiling in a mouse model of cerebrovascular pathology, associated with the two most prevalent forms of dementia, namely AD and VCID, we identified multiple proteins demonstrating significantly altered levels. Level dysregulation in these proteins point to perturbations in brain vessel vasomotricity, remodeling, and inflammation. We further demonstrated that several of the differentially-expressed mouse proteins are (a) expressed in human brain vasculature, and (b) found as cargo proteins in EVs from human plasma. Given the growing popularity of EVs in blood as a novel and minimally invasive biomarker discovery platform for the age-related dementias, several of the proteins identified by can serve as protein biomarkers in humans.

## ACKNOWLEDGEMENTS

We thank Ms. C. Delaney, Mr. L. Tessier, Mr. S. Williamson and Mr. K. Chan (HHT-National Research Council Canada) for their technical assistance with protein isolation and mass spectrometry.

## CONFLICT OF INTEREST

All authors (ASH, ZM, ATS, FD, DBS, EH, AB) declare having no personal or financial competing conflict of interests.

## DISCLOSURE

### Funding

The author(s) disclosed receipt of the following financial support for the research, authorship, and/or publication of this article: Grants (to EH) from Canadian Institutes of Health Research (CIHR-MOP-126001), and (to AB) from the Fonds de Recherche Québec - Santé (FRQS: Chercheur boursiers Junior 1, 2020-2024), and the Fonds de soutien à la recherche pour les neurosciences du vieillissement from the Fondation Courtois. Scholarships (to ZM) from the Vascular Training Platform (VAST, 2023-2024), and (to FD) from Bourse de Mérite de la faculté de Médecine de l’Université de Montréal (2021-2022) and the Fonds de Recherche Québec - Santé (FRQS; Bourse de formation de maîtrise, 2022-2023).

## SENTENCE REGARDING SUPPLEMENTARY INFORMATION

Supplementary material provided.

## Notes

### Competing Interest Statement

The authors have declared no competing interest.

