## Supplemental material for "Proteome profiling of brain vessels in a mouse model of cerebrovascular pathology"

### SUPPLEMENTARY MATERIAL

#### Cerebrovascular proteomics workflow

##### Protein extraction from cerebral arteries

Briefly, 100 μL of 0.2% (w/v) Rapigest SF (Waters, Milford, MA, USA) in 50 mM ammonium bicarbonate was added to each of the three biological replicates and subjected to three, short pulses (20 sec each) of sonication for protein solubilization, followed by incubation (95°C dry bath, 10 min), and centrifugation (20 min, 10,000 g). The supernatant (S0) containing Rapigest SF-solubilized proteins was transferred to a new tube, and 20 μL removed for (a) total protein content analyses, and (b) to run a 1D, 12% SDS-PAGE gel. Conversely, the pellet was resuspended in 50 μL of buffered collagenase (0.1 mg/mL crude collagenase, 50 mM Tris, pH 7.5, 2 mM CaCl2) and incubated (60 min at 37°C) to further solubilize extracellular matrix and membrane proteins. Thereafter, 80 μL of sample-specific S0 was added to the above mix, and the entire mixture sonicated three times (30 sec each), followed by incubation (10 min, 95°C), and centrifugation (20 min, 10,000 g). The final supernatant (S1) containing solubilized cerebral arterial proteins was collected.

##### Protein digest

Proteins in each S1 supernatant were denatured in 100 mM 2-mercaptoethanol (10 min, 95°C). Following a 2 min cool down at room temperature (RT), 6 μL of N-Glycosidase F (Roche, Cat. # 11365185001) was added to each sample and incubated overnight at 37°C to deglycosylate glycoproteins, which increases detection of membrane and secreted proteins by mass spectrometry [1]. Thereafter, each sample was reduced in fresh 4 mM dithiothreitol (10 min at 95°C), and then alkylated in the dark in 10 mM freshly prepared iodoacetic acid (30 min at RT). Subsequently, each sample was diluted to 450 μL with Milli-Q water and 5 μL of trypsin enzyme stock solution (Trypsin Gold, mass spectrometry grade, Promega, Cat. # V5280) added, and then incubated overnight (37°C).

##### Maximizing proteome coverage

As described in detail previously [1,2], we used gel-free proteomics to maximize cerebral arterial proteome coverage. Specifically, tryptic-digest of each S1 protein mix was followed by fractionation of peptides by strong cation exchange [3], and eluted peptide fractions collected from low to high salt-containing cation exchange elution buffer concentration, by slowly injecting (~1 drop/sec) 0.5 ml of each elute buffer. Specifically, peptide fractions were collected at 20%, 40% and 100% elution buffer concentrations [3].

##### Mass spectrometry analysis

For all 18 samples, each strong cation exchange fraction was analyzed by nanoLC-MS/MS, culminating in a grand total of 162 runs. Specifically, each fraction was analyzed using ESI-LTQ-Orbitrap-XL mass spectrometer (Thermo) coupled to a NanoAcquity UPLC system (Waters, Milford, MA), as previously described [1,2].

*Protein identification and bioinformatics*

The raw data was converted to mzXML and MGF (Mascot generic format) format files using Msconvert (<https://proteowizard.sourceforge.io/>). The resulting MS/MS (tandem mass spectrometry) spectra were searched against Mus musculus Swiss-Prot database using Mascot v2.2.0 search engine [4] for protein identification with the following parameters: enzyme = trypsin; modifications = C (carbamidomethyl, fixed), M (oxidation, variable); peptide tolerance = 1.5 Da; fragment tolerance = 1.5 Da; one missed cleavage allowed). Decoy database searches were used to estimate the false positive identification rate of peptides. Removal of peptides with a delta mass >15 ppm or a score <35 yielded a false positive identification rate of 1%. In addition, Peptide Prophet probabilities, an independent statistical measure of peptide identification, yielded *p*≥ 0.95 for all identified peptides. To extract quantitative MS (mass spectrometry) data, to align all runs and integrate protein search results, in-house software MatchRx 2 version QnD-2.0 was used. Peptide intensity validation was performed using MSight software version 1.0 (<http://www.expasy.ch/MSight>).

#### Supplemental Figures

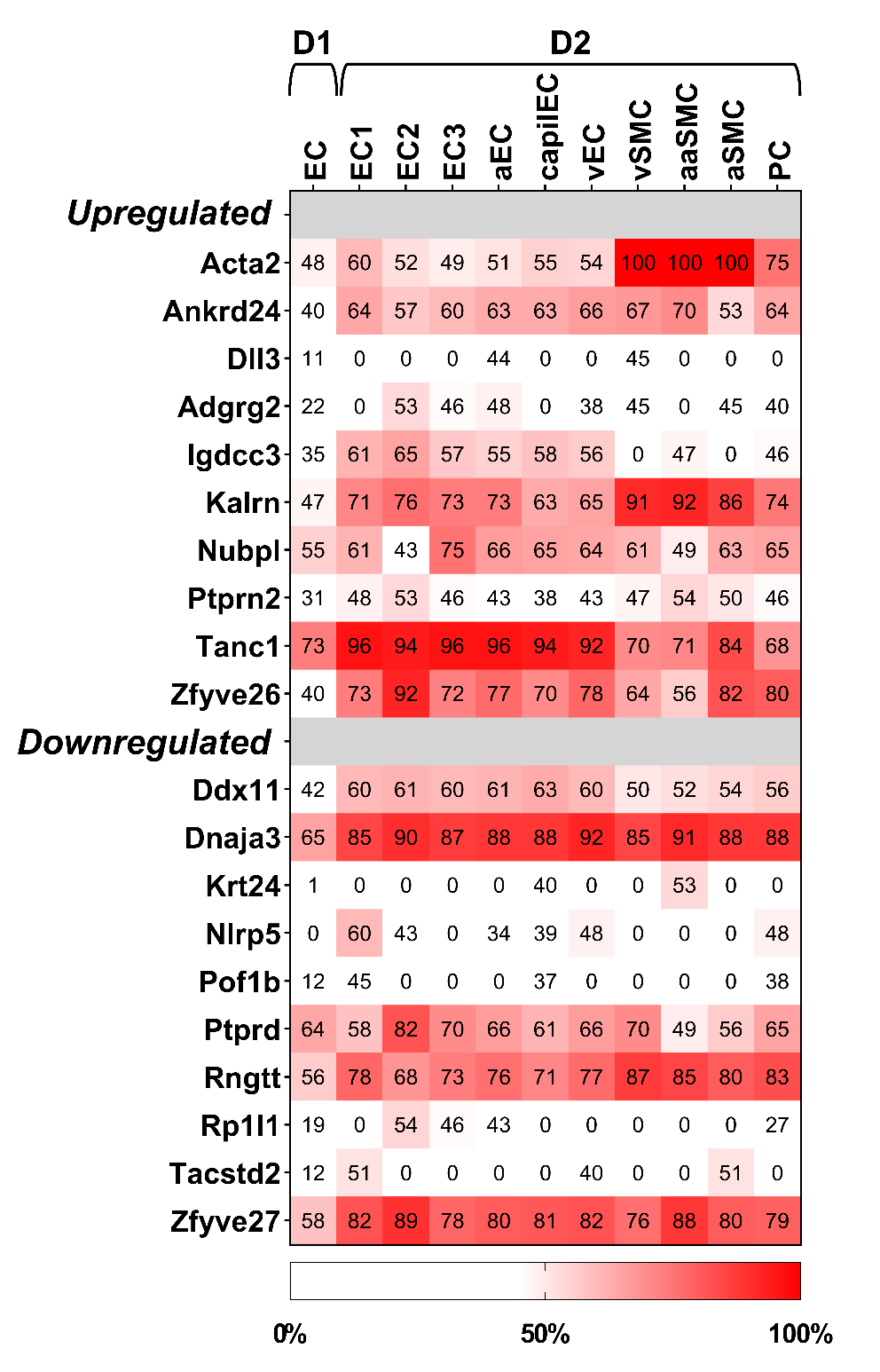

**Supplemental Figure S1**: **Relative gene expression of List 2 proteins in public vascular datasets.** Relative gene expression of the 20 List 2 proteins in mouse cerebrovascular single cell transcriptome databases D1 [5] and D2 [6]. Abbreviations: EC, endothelial cell; aEC, arterial EC; capilEC, capillary EC; vEC, venular EC; vSMC, venular smooth muscle cell; aaSMC, arteriole SMC; aSMC, arterial SMC; PC, pericyte. This figure is duplicated from Figure 4 except raw values are on the heat map.

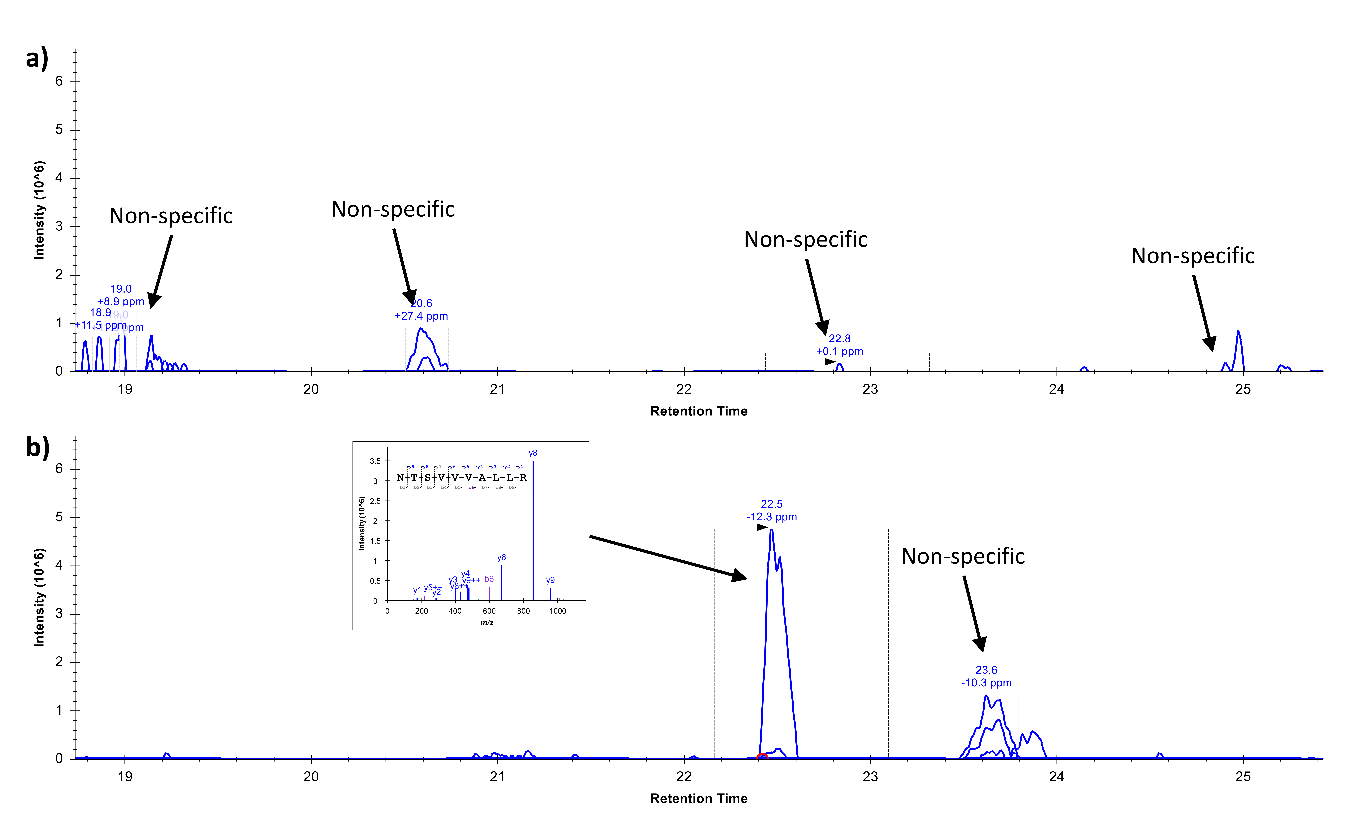

**Supplemental Figure S2**: **Examples of extracted ion-chromatograms and MS/MS spectra in precleared plasma and depleted plasma.** (a and b) An example of extracted ion-chromatogram for a TANC1 peptide in non-depleted (a) and depleted (b) plasma. Arrows indicate either non-specific peaks or specific peak. TANC1-specific peak was only detected in depleted plasma (b) as confirmed by its MS/MS spectra.

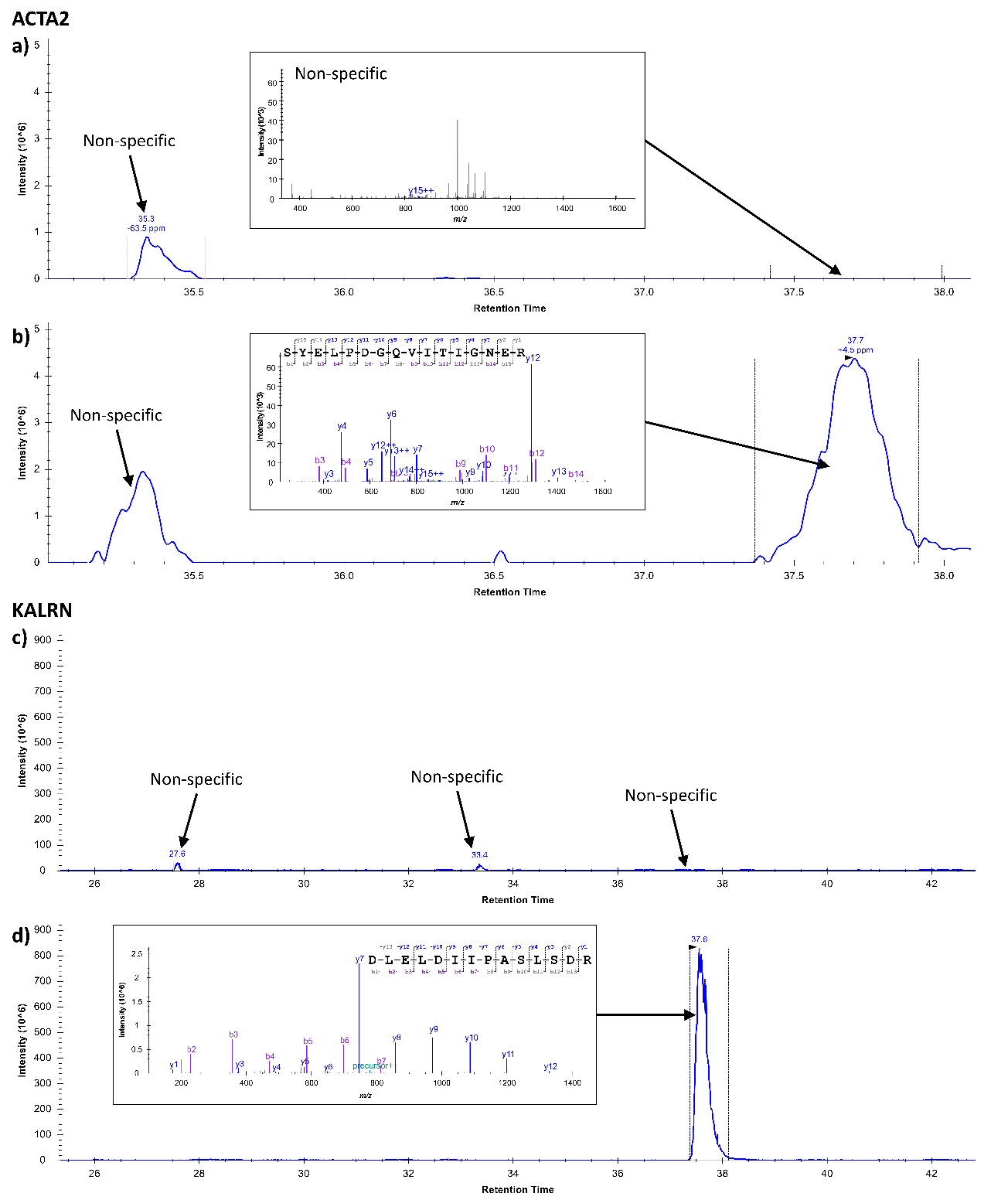

**Supplemental Figure S3**: **Examples of extracted ion-chromatograms and MS/MS spectra in precleared plasma and total plasma EVs.** (a and b) An example of extracted ion-chromatogram for an ACTA2 peptide in non-depleted plasma (a) and total EVs (b). Arrows indicate either non-specific peaks or specific peak. ACTA2-specific peak was only detected in total EVs (b) as confirmed by its MS/MS spectra. (c and d) An example of extracted ion-chromatogram for an KALRN peptide in non-depleted plasma (c) and total EVs (d). Arrows indicate either non-specific peaks or specific peak. KALRN-specific peak was only detected in total EVs (d) as confirmed by its MS/MS spectra.

#### Supplemental Table

| **Uniprot** (ID) | **Gene** | **Gene** (synonyms) | **Protein Name**  (recommended) | **Log2 fold** change | **EVs** | | | |
| --- | --- | --- | --- | --- | --- | --- | --- | --- |
|  |  |  |  |  | **M** | | **H** | |
|  |  |  |  |  | p | r | p | r |
| Q3UHJ0 | **Aak1** | Kiaa1048 | AP2-associated protein kinase 1 | -1.09 | X |  | X | X |
| P62737 | **Acta2** | Actsa, Actvs | Actin, aortic smooth muscle | 1.05 | X |  | X | X |
| Q8CJ12 | **Adgrg2** | Gpr64, Me6 | Adhesion G-protein coupled receptor G2 | 4.69 | X |  | X | X |
| Q80VM7 | **Ankrd24** | - | Ankyrin repeat domain-containing protein 24 | 1.08 |  |  | X |  |
| Q8BVM2 | **Antxrl** | - | Anthrax toxin receptor-like | -1.61 |  |  |  |  |
| O88879 | **Apaf1** | - | Apoptotic protease-activating factor 1 | -1.33 | X |  | X | X |
| Q9Z1K5 | **Arih1** | Ari, Ubch7bp | E3 ubiquitin-protein ligase ARIH1 | -1.3 |  |  | X | X |
| Q80VC9 | **Camsap3** | Kiaa1543 | Calmodulin-regulated spectrin-associated protein 3 | -2.18 |  |  |  | X |
| Q80ZU5 | **Ccdc181** | - | Uncharacterized Coiled-coil domain-containing protein 181 protein C1orf114 homolog | 1.33 |  |  |  |  |
| P06909 | **Cfh** | Hf1 | Complement factor H | 1.14 | X |  | X | X |
| Q99388 | **Csprs** | D1Lub1 | Component of Sp100-rs | -2.05 |  | X |  |  |
| Q02248 | **Ctnnb1** | Catnb | Catenin beta-1 | -1.32 | X |  | X | X |
| A2AKB9 | **Dcaf10** | Wdr32 | DDB1- and CUL4-associated factor 10 | -1.25 |  |  |  | X |
| Q91VR5 | **Ddx1** | - | ATP-dependent RNA helicase DDX1 | 1.11 | X |  | X | X |
| Q6AXC6 | **Ddx11** | - | Probable ATP-dependent RNA helicase DDX11 | -1.08 |  |  | X | X |
| Q9CWT6 | **Ddx28** | - | Probable ATP-dependent RNA helicase DDX28 | 1.35 |  |  | X | X |
| Q9R0Z9 | **Dlc1** | Arhgap7, Stard12 | Rho GTPase-activating protein 7 | 1.71 |  |  | X | X |
| Q811D0 | **Dlg1** | Dlgh1 | Disks large homolog 1 | -1.46 | X |  | X | X |
| O88516 | **Dll3** | - | Delta-like protein 3 | 1.48 |  |  | X | X |
| Q99M87 | **Dnaja3** | Tid1 | DnaJ homolog subfamily A member 3, mitochondrial | -1.95 | X |  | X | X |
| Q8BVG4 | **Dpp9** | - | Dipeptidyl peptidase 9 | 1.62 | X |  | X | X |
| O70251 | **Eef1b** | Eef1b2 | Elongation factor 1-beta | 1.51 | X |  | X | X |
| O08810 | **Eftud2** | Snrp116 | 116 kDa U5 small nuclear ribonucleoprotein component | -2.28 | X |  | X | X |
| Q6DYE8 | **Enpp3** | - | Ectonucleotide pyrophosphatase/phosphodiesterase family member 3 | 1.01 | X |  | X |  |
| Q9D281 | **Fam114a1** | Noxp20 | Protein Noxp20 | -1.54 |  |  | X |  |
| Q8K285 | **Fcho1** | - | F-BAR domain only protein 1 | -1.65 |  |  | X | X |
| Q8CG64 | **Fkrp** | - | Ribitol 5-phosphate transferase FKRP | 1.01 | X |  |  | X |
| Q7TQ32 | **Hjv** | Hfe2, Rgmc | Hemojuvelin | -1.84 |  |  |  |  |
| Q8R0J8 | **Idnk** | - | Probable gluconokinase | 1.03 |  |  | X | X |
| Q8BQC3 | **Igdcc3** | Punc | Immunoglobulin superfamily DCC subclass member 3 | 1.46 |  |  |  | X |
| Q920Q8 | **Ivns1abp** | Kiaa0850, Nd1, Nd1L, Nd1S, Ns1, Ns1bp | Influenza virus NS1A-binding protein homolog | -1.55 |  |  |  | X |
| A2CG49 | **Kalrn** | - | Kalirin | 1.02 |  |  | X | X |
| Q03719 | **Kcnd1** | - | Potassium voltage-gated channel subfamily D member 1 | -1.75 |  |  | X | X |
| P28740 | **Kif2a** | Kif2, Kns2 | Kinesin-like protein KIF2A | -1.12 | X |  | X | X |
| Q8VCK5 | **Klhl20** | Kiaa4210, Kleip | Kelch-like protein 20 | -1.15 |  |  | X | X |
| A1L317 | **Krt24** | Ka24 | Keratin, type I cytoskeletal 24 | -1.02 | X |  | X |  |
| Q0P5X1 | **Lrriq1** | - | Leucine-rich repeat and IQ domain-containing protein 1 | -1.14 | X |  | X |  |
| Q9QYR6 | **Map1a** | Mtap1, Mtap1a | Microtubule-associated protein 1A | -1.74 |  |  | X | X |
| Q9WUU9 | **Mcm3ap** | Ganp, Map80 | Germinal-center associated nuclear protein | -1.06 |  |  | X | X |
| Q6PDC8 | **Mfsd4** |  | Major facilitator superfamily domain-containing protein 4 | 1.78 |  |  | X |  |
| P97479 | **Myo7a** | Myo7 | Unconventional myosin-VIIa | -1.51 | X |  | X |  |
| Q9D5Y0 | **na** | - | Uncharacterized protein C7orf31 homolog | 1.39 |  |  |  |  |
| Q9CQH3 | **Ndufb5** | - | NADH dehydrogenase [ubiquinone] 1 beta subcomplex subunit 5, mitochondrial | -1.4 | X |  | X | X |
| Q9R1M5 | **Nlrp5** | Mater, Nalp5 | NACHT, LRR and PYD domains-containing protein 5 | -1.05 |  |  |  |  |
| Q9CWD8 | **Nubpl** | - | Iron-sulfur protein NUBPL | 2.39 |  |  | X | X |
| Q6P3D0 | **Nudt16** | - | U8 snoRNA-decapping enzyme | -1.32 |  |  | X | X |
| Q61036 | **Pak3** | Pak-3, Pakb, Stk4 | Serine/threonine-protein kinase PAK 3 | -2.25 | X |  | X |  |
| P61249 | **Pde6h** | - | Retinal cone rhodopsin-sensitive cGMP 3',5'-cyclic phosphodiesterase subunit gamma | -1.51 |  |  |  |  |
| Q8BUY9 | **Pggt1b** | - | Geranylgeranyl transferase type-1 subunit beta | -1.07 |  |  | X | X |
| P26450 | **Pik3r1** | - | Phosphatidylinositol 3-kinase regulatory subunit alpha | -1.09 |  |  | X | X |
| Q8K4L4 | **Pof1b** | - | Protein POF1B | -1.76 |  |  | X |  |
| Q8R4S0 | **Ppp1r14c** | Kepi | Protein phosphatase 1 regulatory subunit 14C | 1.27 |  |  |  | X |
| Q8BQ30 | **Ppp1r18** | - | Phostensin | 1.17 |  | X | X | X |
| Q0VGB7 | **Ppp4r2** | - | Serine/threonine-protein phosphatase 4 regulatory subunit 2 | -1.07 |  |  | X | X |
| Q91WG5 | **Prkag2** | - | 5'-AMP-activated protein kinase subunit gamma-2 | 1.2 | X |  | X | X |
| Q64487 | **Ptprd** |  | Receptor-type tyrosine-protein phosphatase delta | -1.45 | X |  | X |  |
| P80560 | **Ptprn2** | - | Receptor-type tyrosine-protein phosphatase N2 | 1.23 |  |  |  | X |
| O55236 | **Rngtt** | Cap1a | mRNA-capping enzyme | -1.41 |  |  | X | X |
| Q8CGM2 | **Rp1l1** | Rp1hl1 | Retinitis pigmentosa 1-like 1 protein | -1.34 |  |  |  |  |
| O70258 | **Sgce** | - | Epsilon-sarcoglycan | -1.97 | X |  | X | X |
| Q80W37 | **Snupn** | Rnut1 | Snurportin-1 | -1.02 |  |  | X | X |
| Q505B8 | **Syce2** | Cesc1 | Synaptonemal complex central element protein 2 | 1.07 |  |  |  |  |
| Q9CUU3 | **Sycp2** | Scp2 | Synaptonemal complex protein 2 | -1.21 | X |  | X |  |
| Q8BGV3 | **Tacstd2** | Trop2 | Tumor-associated calcium signal transducer 2 | -1.31 |  |  | X |  |
| Q0VGY8 | **Tanc1** | - | Protein TANC1 | 1.11 | X |  | X | X |
| Q7TQA6 | **Tas2r38** | T2r31, Tas2r138 | Taste receptor type 2 member 38 | 1.31 |  |  |  |  |
| P70325 | **Tbx4** | - | T-box transcription factor TBX4 | 1.63 |  |  |  |  |
| A8C756 | **Thada** | Kiaa1767 | Thyroid adenoma-associated protein homolog | 2.62 |  |  | X | X |
| P39447 | **Tjp1** | Zo1 | Tight junction protein ZO-1 | -1.12 | X |  | X | X |
| Q6A070 | **Togaram1** | Fam179b, Kiaa0423 | TOG array regulator of axonemal microtubules protein 1 | 1.39 |  |  |  |  |
| Q9CYZ2 | **Tpd52l2** | - | Tumor protein D54 | -1.09 | X |  | X | X |
| Q80VC6 | **Trnau1ap** | Secp43, Trspap1 | tRNA selenocysteine 1-associated protein 1 | 1.53 |  |  |  | X |
| Q8VCH8 | **Ubxn4** | Ubxd2, Ubxdc1 | UBX domain-containing protein 4 | 2.33 |  |  | X | X |
| Q9JKB1 | **Uchl3** | - | Ubiquitin carboxyl-terminal hydrolase isozyme L3 | -2.1 | X |  | X | X |
| Q91YD6 | **Vill** | Villp | Villin-like protein | -1.11 |  |  | X | X |
| Q8BYK8 | **Zc3h6** | Zc3hdc6 | Zinc finger CCCH domain-containing protein 6 | -1.12 |  |  | X |  |
| Q5DU37 | **Zfyve26** | Kiaa0321 | Zinc finger FYVE domain-containing protein 26 | 1.13 |  |  | X | X |
| Q3TXX3 | **Zfyve27** | - | Protrudin | -1.56 |  |  | X | X |

**Supplemental Table 1: List 1 and List 2 proteins.** List 2 proteins are highlighted in yellow. Note that the Vesiclepedia database, was used to identify List 1 and 2 proteins detected in the TGF cerebrovasculature, and also detected in human extracellular vesicles. Abbreviations: na, not available; EVs, extracellular vesicles; M, mouse; H, human; p, protein; r, mRNA.
